# Encoding multiple virtual signals in DNA barcodes with single-molecule FRET

**DOI:** 10.1101/2020.11.12.379479

**Authors:** Sung Hyun Kim, Hyunwoo Kim, Hawoong Jeong, Tae-Young Yoon

## Abstract

DNA barcoding provides a way to label a huge number of different biological molecules using the extreme programmability in DNA sequence synthesis. Fluorescence imaging is an easy-to-access method to detect individual DNA barcodes, which can be scaled up to a massively high-throughput format. Large overlaps between emission spectra of fluorescence dyes, however, severely limit the numbers of DNA barcodes–and thus its signal space–that can be detected in a simultaneous manner. We here demonstrate the use of single-molecule fluorescence resonance energy transfer (FRET) to encode virtual signals in DNA barcodes using conventional two-color fluorescence microscopy. By optimizing imaging and biochemistry conditions for weak hybridization events for DNA barcodes, we markedly enhanced accuracy in our determination of the efficiency by which single-molecule FRET occurred. This allowed us to unambiguously differentiate six DNA barcodes exhibiting different FRET values without involving probe sequence exchanges. Our method can be directly incorporated with previous DNA barcode techniques, and may thus be widely adopted to expand the signal space of the DNA barcode techniques.

---

DNA, the basic material chosen by biological systems to carry and preserve their genetic information, shows a remarkable level of chemical stability and mechanical rigidity^1–3^. DNA shows high specificity in its hybridization to make the helical double-stranded structure. Insertion of one mismatch in otherwise perfectly matching sequences of 20 nucleotides reportedly weakens the hybridization strength up to ten folds^4, 5^.

This high specificity in DNA hybridization finds its exquisite use in many different areas of DNA nanotechnologies, with one prominent application being programmable labeling of different biological molecules for optical detection^6, 7^. Compared with other labeling and detection methods, such as those harnessing isotopes^8^ and metabolites^9^, fluorescence labeling and optical imaging is by far an easy-to-access and high-throughput method for identification of biological molecules^10, 11^. Harnessing the extreme programmability of DNAs, it has been shown that myriads of biological molecules are labeled with different sequences of DNAs and detected with the high specificity of DNA hybridization. These technologies, collectively termed DNA barcoding, open a way to cope with the ever-increasing complexity of biological molecules and investigate their spatiotemporal distributions up to the single-cell level^12–15^.

Relatively broad emission spectra of fluorescent dyes, however, make it already challenging to detect more than three fluorescent dyes simultaneously^16^. Several technologies have been reported to increase either the signal space spanned by DNA barcodes or the accuracy in the detection of successful DNA hybridization events. To increase the signal space of DNA barcodes, different sets of DNA probes are introduced in a sequential manner and binding patterns observed through these multiple rounds of probe introduction collectively encode unique identities of individual biological molecules^17–19^. For example, MERFISH, which identifies individual DNA barcodes from two-digit binding patterns of different DNA probes, has demonstrated a detailed mapping of more than thousands of genomic RNAs in small cellular nuclei^20^. DNA point accumulation for imaging in nanoscale topography (DNA-PAINT) uses deliberately weak hybridization of DNA probes to induce recurrent binding of DNA probes. A marked increased accuracy in the detection of DNA hybridization has been demonstrated because the recurrent binding of DNA-PAINT probes increased the number of fluorescent photons detected for individual targets^21^.

In addition, single-molecule fluorescence resonance energy transfer (FRET) is finding ways to be integrated with DNA barcoding technologies^22–24^. For example, single-molecule FRET is shown to decrease the fluorescence background noise coming from diffusing DNA probes, thereby substantially increasing the detection efficiency^25^. Depending on the distance between donor and acceptor dyes, single-molecule FRET can occur via different efficiencies, and it has been demonstrated that individual DNA hybridization events can be labeled by low and high FRET efficiencies, a remarkable invention that increases the signal depth of DNA barcoding by a factor of two^26^. However, the relatively low signal-to-noise ratios obtained thus far have deterred a more-wide use of the virtual signal channels generated by single-molecule FRET in the DNA barcoding field.

In this work, we report systematic ways to enhance the signal-to-noise ratio in labeling DNA barcodes with different single-molecule FRET efficiencies. As a result, we were able to demonstrate dissection of the FRET efficiency space into six virtual channels. Given that the previous state-of-art demonstration that simply divided the FRET space into low- and high-FRET states, our results represent a further expansion of the signal space of the DNA barcode technology by a factor of three. We note the current expansion in the signal space can be readily combined with other DNA barcoding technologies without requiring complicated instrumentation.

The key design principle for our creating multiple virtual FRET channels is to detect individual FRET barcodes in a time-differentiated manner. To this end, we deliberately weakened the interaction between barcode and probe sequences. These weak interactions have been shown to render single hybridization events largely transient, yielding chances for other barcode sequences to be searched by probe sequences. This allows the manifestation of different FRET efficiencies encoded by the individual barcodes in the time domain. We thus deemed accurate FRET detection during transient hybridization events as the first step to achieving our goal.

To examine this possibility, we constructed a hybrid DNA structure, in which a FRET barcode was encoded in a single-stranded (ss) overhang (**Fig. 1a**). This barcode sequence contains binding sites for two short stretches of ssDNAs labeled with Cy3- and Cy5 dyes each (hereafter referred to as cy3- and cy5-probe) such that when bound with both probes, FRET is induced with a predesigned efficiency (*E*) (**Fig. 1a** and **Supp. Fig 1a**). To ensure transient binding of the probes, we limited the lengths of the binding sequences to 8 and 7 nucleotides (nt) for the cy3- and cy5-probes, respectively. The sequence of the barcodes and probes are adapted from the consensus sequences used for DNA-PAINT and modified by trimming the end nucleotides for faster binding kinetics (**Supp. Fig. 1a**)^18^. After assembly of the hybrid DNA structures on the polymer-passivated surface of a microfluidic chamber^27^, we added both the cy3- and cy5-probes to the chamber (each at 100 nM and 500 nM, respectively) and took time-resolved fluorescence imaging at the single-molecule level (a time resolution of 100 ms) (**Fig. 1b** and **Supp. Fig. 2**). We reasoned that recurrent binding and unbinding of the cy3 and cy5 probes would give rise to stepwise intensity fluctuations within certain spatial regions. After accumulating images from our time-resolved recording, we thus searched for regions with large standard deviations in their fluorescence signals and designated these regions as regions of interests (ROIs) (**Fig. 1b-c**).

**Figure 1.**
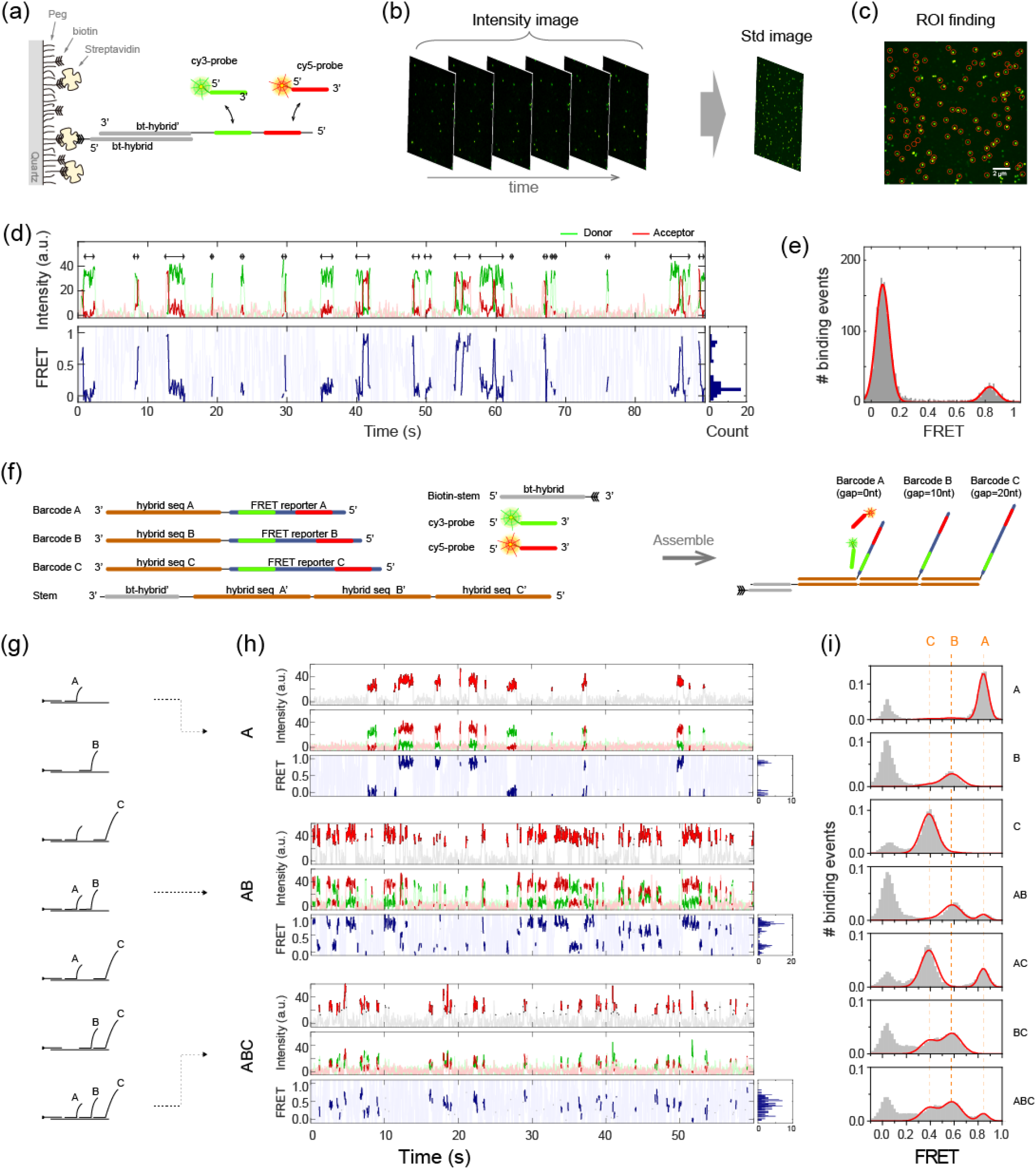
Time-differentiated single-molecule FRET barcode detection. (a) Schematics of the hybrid DNA complexes for transient FRET detection. (b-c) An example time-series of single-molecule fluorescence images and the intensity std image for fluorescence ROI finding. ROI are highlighted with a red circle. (d) A representative fluorescence intensity (upper panel) and FRET (bottom panel) time traces. The high-intensity data points for individual binding events were highlighted with double-headed arrows and dark colors. (e) FRET histograms from the probe binding events from all ROIs in the single field of view. (f) Schematics of the hybrid DNA complexes with three FRET barcodes. (g) Seven different combinations of the DNA hybrid structure with three FRET barcodes. (h) Example single-molecule fluorescence and FRET time traces obtained from the hybrid DNA complex consisting of (1st row) barcode [A], (2nd row) barcodes [A,B] and (3rd row) barcodes [A,B,C]. The top panels show the intensity sum of the donor and acceptors, which are separately shown in the mid panels. The binding events were highlighted with dark colors. The bottom panels show the FRET efficiencies calculated. (i) FRET efficiency distribution of the peaks collected from a single field of view for the seven different DNA hybrids. Solid lines are multi-Gaussian fits. Dashed lines are eye-guides for the position of each sub-population.

Time traces extracted from individual ROIs indeed showed recurring, stepwise changes in both cy3 and cy5 fluorescence channels due to the binding of the corresponding probes (**Fig. 1d, upper panel**). We further applied an intensity threshold found by 2-state K-means clustering to identify segments in individual time-resolved traces that exhibit proper fluorescence intensities, indicative of binding of single cy3- and cy5-probes onto the barcode sequence (**Fig. 1d, double-headed arrows**). As expected, we observed a high FRET efficiency (*E*~0.8) when both cy3- and cy5-probes were simultaneously bound, which was reconfirmed in the FRET efficiency distribution histogram built from total 2445 binding events from 176 ROIs (**Fig. 1e**). Binding of only cy3-probes appeared as another peak at *E*~0.1.

As we successfully implemented methods to resolve FRET signals from the transient binding events lasting only a few seconds, we proceeded to resolve multiple barcode sequences with different FRET values. We constructed a similar DNA hybrid structure that had double-stranded stem regions with protruded ssDNA overhangs that contained barcode sequences (**Fig. 1f and Supp. Fig. 1b**). In particular, we varied the spacer length–consisting of poly(thymine)–between the cy3- and cy5-probe binding sites to encode three different FRET barcodes.

We prepared seven hybrid DNA structures with various combinations of the FRET-barcode sequences (**Fig. 1g**). Following the method we developed above with the case of a single FRET barcode, we identified ROIs and transient hybridization events in individual time traces (**Fig. 1h** and **Supp. Fig. 3**). We first examined a hybrid structure with barcodes A and B.

Individual time-resolved trace obtained from this structure reflected two main FRET populations with one peak positioned in the high FRET region (*E*~0.85) and the other in the middle (*E*~0.6), which agreed well with the short and intermediate spacer lengths of 2 and 10 nt used for barcodes A and B, respectively. Likewise, individual traces obtained from a hybrid structure with all three barcodes (A, B, and C) showed a FRET distribution that was fitted well into three Gaussian distributions. Finally, when we built collective FRET histograms for all seven different hybrid structures, we obtained distinct FRET distributions that unambiguously reflected the existence of specific barcode sequences (**Fig. 1i**). Together, these results demonstrate that using only two physical fluorescence channels, we were able to detect multiple FRET barcodes within a diffraction-limited volume, particularly, in a one-pot reaction scheme that does not require buffer exchange for different probe sequences.

Having demonstrated successful detection of the three FRET barcodes, we sought to further increase the detection capacity of our method. We found that the large overlap between the FRET peaks (**Fig. 1i**) offered a practical challenge for increasing the number of barcodes that could be identified in a single measurement. To reduce the uncertainty in the determination of the FRET efficiency for each barcode sequence, we systematically examined several physical factors in our single-molecule imaging. Firstly, we noticed that some ROIs did not show stepwise fluorescence intensity changes expected for single probe association and dissociation, but rather arbitrary intensity fluctuations or large background noise that led to increased noise in our FRET determination **(Fig. 2a-b).** We reasoned that single probe association and dissociation, described by the first-order kinetics, leads to larger dynamic deviation and reduced auto-correlation in the time domain than exhibited by irregular intensity fluctuation of junk fluorescence molecules. Indeed, we found that the barcode traces were clearly distinguishable with their characteristic kinetic time scales which can be quantified by Allan deviation and auto-correlation values (**Fig. 2c-d**). With a proper selection of lag times (around 1 s) that separates the two groups maximally, we were able to achieve ~99% true-positive and ~97% false-negative detection rates (**Fig. 2e**).

**Figure 2.**
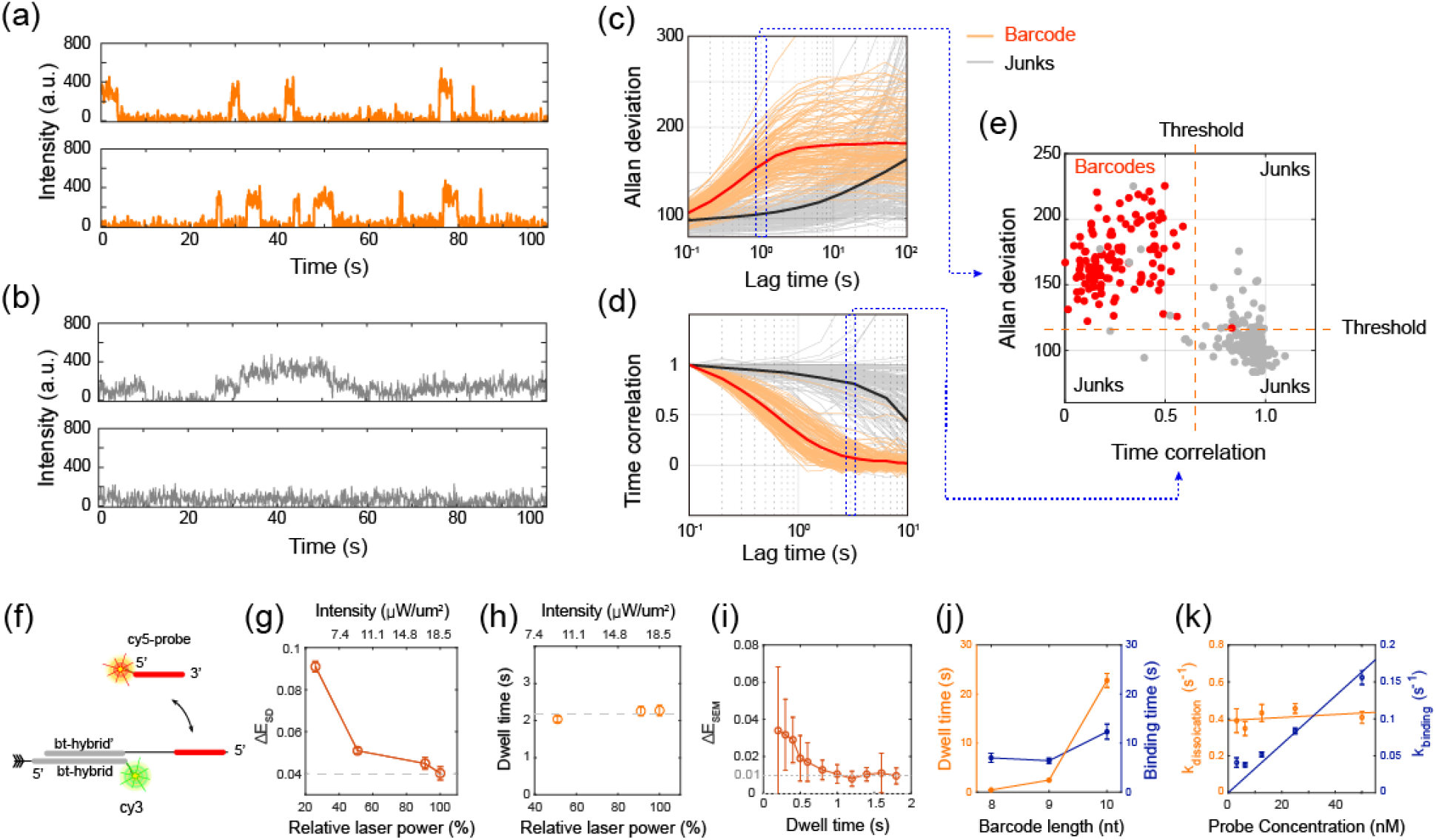
Experimental and post-experimental optimization of the FRET barcode detection accuracy. (a, b) Example traces of (a) the single-molecule barcode signals and (b) the junk signals. (c-d) Discrimination of the junk signals from the barcode signals by checking the time-correlation and Allan deviation. (c) Allan deviations and (d) the time-correlations for individual ROI were calculated for varying lag times. Individual ROI were manually classified into barcode (orange lines) and junks (grey lines). Thick lines are the averaged Allan deviations and time correlations for all the barcodes (red) or junks (black). (e) A representative 2D correlation plot for the time-correlation and Allan deviations for selected lag times that maximally separate the barcodes and junks. Orange dashed lines define the thresholds to discriminate the barcodes from junks. (f) Schematic of the DNA hybrid used for the barcode detection optimization in (g-k) (g) FRET errors (SD) measured at various excitation laser intensities. Gray dotted line is an eye-guide to the asymptotic limit of 0.04. (h) Dwell times of the cy5-probe at various laser powers. Gray dotted line is an eye-guide for horizontal comparison. (i) Barcode determination accuracy (SEM) calculated for various probe dwell times. (j) Probe length dependence on dwell times and binding times (average time spent for a new binding event from the previous probe dissociation). (k) Probe concentration dependence on the binding kinetics.

After filtering out the junk traces, we sought a way to increase the signal-to-noise ratio of the barcode FRET signals. Our total internal reflection microscopy equipped with an electron-multiplying charge-coupled device (EMCCD) was operated at its maximum photon detection sensitivity, and we examined the standard deviation in the determined FRET efficiencies as we increased the excitation power. For straightforward data acquisition and analysis, we covalently attached a cy3 dye to a hybrid DNA structure and retained a cy5-probe binding site in the ssDNA overhang (**Fig. 2f** and **Supp. Fig. 1c**). The standard deviation (SD) in our FRET efficiency determination (for each frame of 100 ms) decreased with the laser intensity used, approaching an asymptotic limit of 0.04 over ~18.5 μW/μm^2^ (**Fig. 2g**). This persistent standard deviation likely arose from the electronic shot noise in EMCCD. We did not observe a reduction in the dwell time of single binding events, indicating that photobleaching did not affect our determination of dissociation kinetics of the fluorescent probes even at the maximum power used (**Fig. 2h**).

We noted that the accuracy in our FRET barcode determination can be further enhanced beyond the asymptotic limit by averaging the FRET values while single probe binding events last because the accuracy becomes limited by the standard error of the mean (SEM) rather than by SD. When examining this SEM of FRET values with respect to different averaging times, we found the accuracy in our barcode determination further reduced and reached a lower limit of *ΔE*_SEM_~0.01 when the averaging time was longer than 1 s (**Fig. 2i**). We then varied the length of the probe sequence to optimize the dwell time of single binding events (**Fig. 2j**). While probe sequence of 8 nt yielded an average binding dwell time on the order of hundreds of milliseconds, which was presumably too short for reliable FRET determination. A 10 nt-long probe used in our study showed a dwell time of ~20 s, too long that would increase the chances of multiple probe binding within an optical diffraction limit. On the other hand, 9 nt-long cy5-probe returned dwells time of ~2 s, fitting into a desired range that we could use to design a detection system with higher multiplexity. Finally, we investigated the time gap between two successive binding events as a function of the probe concentration, and found that 5 to 50 nM concentrations gave a binding latency between 5 to 20 s (**Fig. 2k**). This binding latency is slightly longer than the binding dwell times determined above and thus separates individual binding events with a reasonable probability, suggesting that the concentration range of 5 to 50 nM might be used with 9 nt-long probes.

After optimizing parameters for a better resolution in discerning individual FRET barcodes, we designed an extensive hybrid DNA structure that could hold up to six FRET barcodes (**Fig 3a** and **Supp. Fig. 1d**). The stem part works as the backbone of the hybrid structure, made up of three ssDNAs–referred to as STEM 1, STEM 2, and Biotinylated STEM–that hybridize with one another. In particular, STEM 1 and STEM 2 carried orthogonal hybridization sequences for adopting three different FRET barcode sequences each.

**Figure 3.**
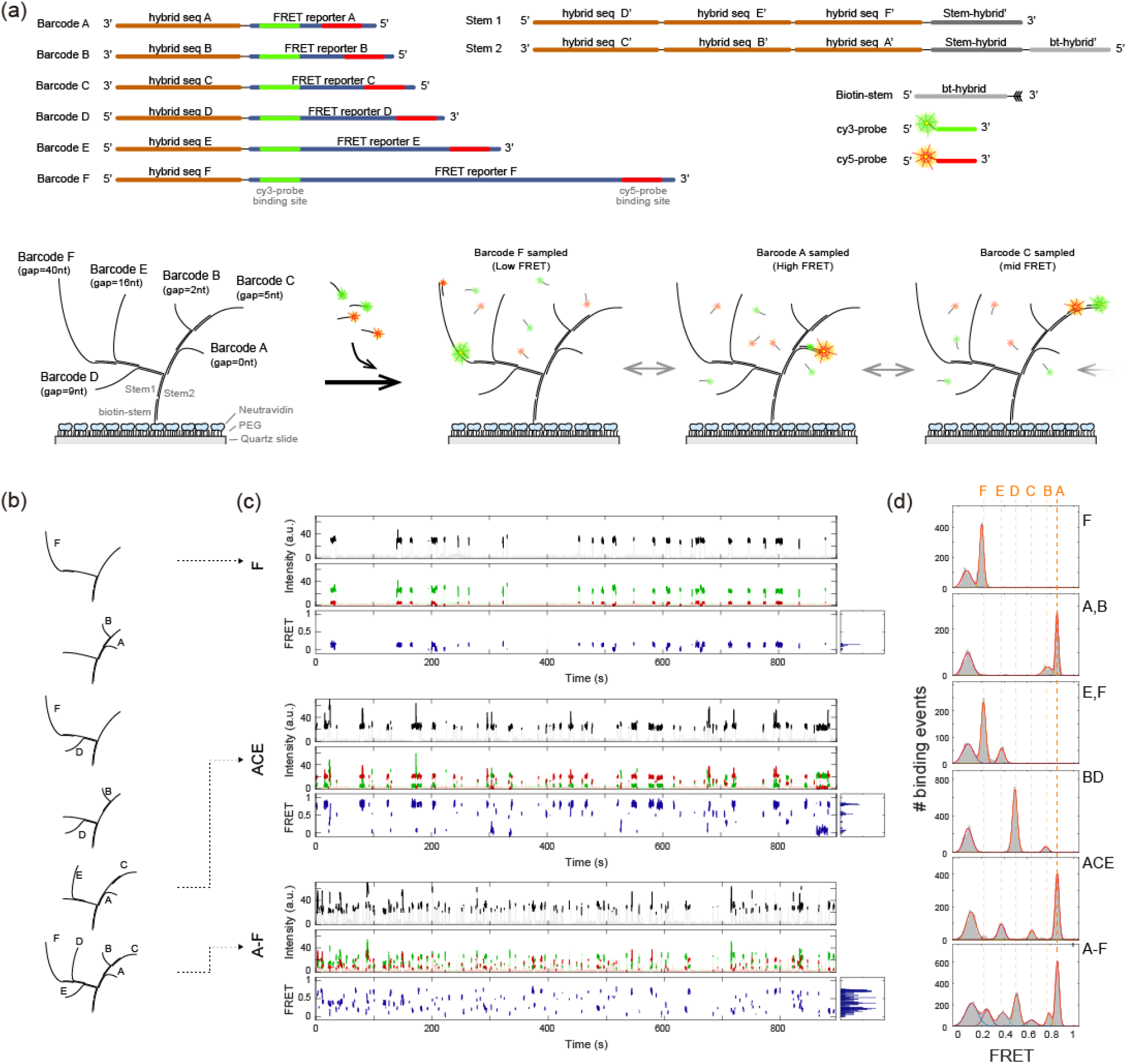
Single-molecule FRET barcode assay and identification of individual components from artificial DNA complexes. (a) Schematics of the hybrid DNA complexes with six FRET barcodes. (b) Depiction of various DNA hybrid structures with different FRET barcode combinations. (c) Example single-molecule fluorescence and FRET time traces obtained from the hybrid DNA complex consisting of barcode [F], barcodes [A,C,E] and barcodes [A,B,C,D,E,F]. The top panels show the intensity sum of the donor and acceptors, which are separately shown in the mid panels. The binding events were highlighted with dark colors. The bottom panels show the FRET efficiencies calculated. (d) FRET efficiency distribution of the peaks collected from a single field of view. From the top, DNA complexes consisting of [F], [A,B], [D,F], [B,D], [A,C,E], and [A,B,C,D,E,F]. Solid lines are Gaussian fits to each sub-population. Dashed lines are eye-guides for the position of each sub-population.

Out of 2^6^ possible combinations of the six barcodes, we attempted to assemble six structures as precedents (**Fig. 3b**). After immobilizing assembled structures on the surface, we added cy3- and cy5-probes to initiate the FRET efficiency sampling. Referring to the binding kinetics determined above, we chose 9 nt-long sequences and 10 and 100 nM concentrations for the cy3 and cy5-probes, respectively. These reaction conditions rendered virtually all barcode sequences fully occupied with the cy5-probes while cy3-probes sampled individual FRET barcode sequence on the order of a few seconds.

When we collected FRET values sampled by the cy3 probes for single hybrid structures, the FRET histogram mirrored the existence of all FRET barcodes present (**Fig. 3c** and **Supp. Fig. 4**). For example, the FRET histogram obtained from a single hybrid structure with barcode F-with the largest spacer sequence of 40 nt–showed a low FRET population at *E*~0.2 beside the donor-only peak at *E*~0.1 (**Fig. 3c**, first row). This FRET pattern was well reproduced when we sampled 225 hybrid structures with the same barcode composition (**Fig. 3d**, first row). In particular, we noted that the FRET peak representing barcode F had a narrow broadness of 0.017, approaching the lower limit of *ΔE*_SEM_~0.01 we found before. For another structure with three barcodes A, C, and E, we observed three clear high FRET populations at E~0.37, 0.64, and 0.86 (**Fig. 3d**, 5^th^ row). Compared with our previous detection of three barcodes in Figure 1J, we noticed that the refined imaging and averaging methods established in Figure 2 enabled far clear resolution of three barcodes in the virtual FRET space. Finally, we formed the hybrid structure incorporating all six FRET barcodes and were able to distinguish all six peaks in the FRET histograms built from 343 structures (**Fig. 3d**, 6^th^ row). This is remarkable because we did not use any buffer exchange for probe exchanges, suggesting that the signal space has been indeed expanded to six channels while using two physical fluorescence channels.

In this report, we demonstrated a DNA barcode method that enables multiplexed detection by creating virtual FRET channels. As a proof-of-concept demonstration, we designed a hybrid DNA structure that contains 6 FRET barcodes within a dimension of ~10 nm. We substantially increased the number of FRET barcodes that can be resolved in a single measurement through optimization of the signal processing and biochemical conditions for probe sampling. Because FRET occurs only when the donor and acceptor dyes simultaneously bind to the target, our FRET-based barcode is much more robust against false positive detection and crosstalk between physical fluorescence channels than typical multi-color fluorescence detection. We envision that addition of one more spectral channel, which creates two orthogonal FRET spaces, the multiplexing capacity would be further increased up to 6^2^ = 36 channels. We note that our multiplexed FRET barcode measurement can be integrated with another recently-developed high-resolution FRET method^28^, which allows accurate FRET determination beyond the classical measurement noise limit from multiple FRET pairs. While multiplexed assay using the DNA barcoding and DNA PAINT methods have been previously demonstrated, these methods rely on introducing different sets of fluorescence probes via microfluidic buffer exchange. Our FRET-barcode method expands the signal space for a given probe set without requiring major changes in instrumentation. We thus expect that our method can be directly incorporated with pre-existing DNA barcode and PAINT methods^18, 19, 24^, providing an avenue toward highly multiplexed imaging of biological information with an increased throughput.

## Materials and Methods

### DNA

The custom-synthesized DNA oligos without chemical modifications were purchased from Macrogen. DNA oligos with amino group, Cy3, Cy5, or biotin modifications were purchased from IDTDNA. The sequences of the barcode and hybrids were adapted from the literature^18^ and modified to avoid stable secondary structure formation. To make the DNA hybrids used in **Fig. 1**, oligos are mixed together with the ratio of 1:2:4:8 for biotin-DNA:STEM 1:STEM 2: FRET-barcodes. The mixture was incubated for 24 hours at room temperature and used without further purification.

### Single-molecule FRET setup

A total internal reflection (TIR) microscope was custom-built around a commercial inverted microscope (IX71, Olympus) equipped with a high-power objective lens (60X, NA1.2, Water immersion, Olympus) and an emCCD (Ixon DU-897, Andor) (**Supp. Fig. 2a**) ^22, 23^. Immobilized samples on a quartz slide glass were illuminated by a 532nm laser and the fluorescence signal was spectrally divided into Cy3 and Cy5 fluorescent channels by using Dual-view (DV-2, Photometrics) before imaged on the emCCD for real-time recording.

### Data analysis

An automated fluorescence image analysis was performed with a home-build image processor and fluorescence time trace analyzer written in Matlab (MathWorks). The analysis codes are freely available from Github (https://github.com/kahutia/transient_FRET_analyzer/releases/tag/v1.3 and https://github.com/kahutia/SingleMoleculeImageAnalyzer/releases/tag/V8.0). First, the initial 1000 frames of the fluorescence images were used to build an intensity standard deviation image. The standard deviation image was then analyzed by a peak finding algorithm to find ROIs. The intensity time traces at each ROI was extracted by taking a Gaussian-weighted sum of the 9 nearest neighbors for every frame. Because the binding of the fluorescence-probes would give a distinct kinetic signature, the traces that show abnormal behavior in time correlation and Allan deviation were regarded as junks and excluded for further analysis. The sums of the donor and acceptor intensities from each time trace were then classified into two groups (high and low-intensity groups) by using 2-state K-means clustering. The average of the low-intensity groups was set to a local intensity background. The high-intensity groups were then further classified by selecting those showing the expected intensity ranges of a single donor-acceptor pair. The selected proper intensity group was then used to calculate the FRET efficiency of each binding event.

## Notes

The authors declare no competing financial interest.

## Acknowledgments

We thank Changwon Kim for active discussions concerning single-molecule FRET imaging. This research was supported by the Bio & Medical Technology Development Program of the National Research Foundation (NRF) funded by the Bio & Medical Technology Development Program (NRF-2018M3A9E2023523). Also, this research was supported by a grant from the National Research Foundation of Korea funded by the Korean government (MSIT) (NRF-2020R1A5A1018081).

## Abbreviations

FRET: fluorescence resonance energy transfer
DNA: Deoxyribonucleic acids
TIRF: microscopy, Total internal reflection fluorescence microscopy.

