## Supplemental information for "Encoding multiple virtual signals in DNA barcodes with single-molecule FRET"

Supplemental Figure 1-4

**Supplemental Figures**

**Supplemental Figure 1.** The sequences of the oligos used for the single-molecule FRET barcode experiments.

(a) DNA sequences used for the single-molecule FRET detection from the transiently binding probes (Fig. 1a-e).

(b) DNA sequences used for the hybrid DNA structures with three FRET barcodes (Fig. 1f-j).

(c) DNA sequences used for the cy3-fixed partial duplex DNA for the barcode detection optimization (Fig. 2f-k).

(d) DNA sequences used for the hybrid DNA structures with six FRET barcodes (Fig. 3).

**Supplemental Figure 2.** Schematics of the experimental setups and the hybrid DNA sample design

(a) Schematic of the prism-type total internal reflection microscopy with two fluorescence detection channels.

(b) Schematic of the sample chamber for the barcoded complex immobilization. The sample chamber is made by sandwiching a quart glass slide and a cover slip via double-sticky tape. The double-sticky tape was pre-cut to make a thin flow chamber and two holes on the quart slide serve as inlet and outlet of the samples.

**Supplemental Figure 3.** Example traces of the DNA hybrids shown in Fig. 1j.

**Supplemental Figure 4.** Example traces of the DNA hybrids shown in Fig. 3d.

Supp. Fig. 1

(a)

(Biotin DNA) 5′-GCCTCGCTGCCGTCG cca-biotin

(Stem-Paired target) CGACGGCAGCGAGGC tt TCTTCATT t GATCTAC

(cy3-probe) 5′Cy3-ctAATGAAGA

(cy5-probe) 5′Cy5-tatGTAGATC

(b)

FRET reporter hybridization sequence

(Barcode A) 5'-TCTTCATT tt GATCTAC tt ATATAGTTTCGTATA
FRET reporter A hybridization sequence A

(Barcode B) 5'-TCTTCATT ttttt ttttt GATCTAC tt CGTCAGTTTACGATT
FRET reporter B hybridization sequence B

(Barcode C) 5'-TCTTCATT ttttt ttttt ttttt ttttt GATCTAC tt TATATGTGTACTATT
FRET reporter C hybridization sequence C

(Stem) 5'-CGACGGCAGCGAGGC cc TATACGAAACTATAT cc AATCGTAAACTGACG CC AATAGTACACATATA
bt-hybrid′ hybridization sequence A′ hybridization sequence B′ hybridization sequence C′

(Biotin-Stem) 5′-GCCTCGCTGCCGTCG cca-biotin
bt-hybrid

(cy3-probe) 5′Cy3-ctAATGAAGA

(cy5-probe) 5′Cy5-tatGTAGATC

(c)

(Stem-target-10nt) TGGCGACGGCAGCGAGGC ttttttttt GATCTACATT

(Stem-target-9nt) TGGCGACGGCAGCGAGGC ttttttttt cATCTACATT

(Stem-target-8nt) TGGCGACGGCAGCGAGGC ttttttttt ctTCTACATT

(Biotin DNA) /5Cy3/GCCTCGCTGCCGTCGCCA-biotin

(cy5-probe) 5′Cy5-AATGTAGATC

(d)

FRET reporter hybridization sequence

(Barcode A) 5′-TCTTCATTA GATCTACAT tt GTAGATTCATGTTTACGATTGTTTCGTATA
FRET reporter A hybridization sequence A

(Barcode B) 5′-TCTTCATTA tt GATCTACAT tt CCATTAACATGTGTACTATTGTTAAGGAAA
FRET reporter B hybridization sequence B

(Barcode C) 5′-TCTTCATTA ttttt GATCTACAT tt CAACAAACTAGTATATCACAGTATGACTTT
FRET reporter C hybridization sequence C

(Barcode D) 5′-CCATGATTATGATTTATCCAGCAAGATTAA t TCTTCATTA ttttt tttt GATCTACAT t
hybridization sequence D FRET reporter D

(Barcode E) 5′-CGAGTTATATGATAAGATAGGATACACATA t TCTTCATTA ttttt ttttt ttttt t GATCTACAT t
hybridization sequence E FRET reporter E

(Barcode F) 5′-CGGTATAATTGACACTAAATGAGAACATAA t TCTTCATTA ttttt ttttt ttttt ttttt ttttt ttttt ttttt ttttt GATCTACAT t
hybridization sequence F FRET reporter F

(Stem 1) 5′-TTAATCTTGCTGGATAAATCATAATCATGG TATGTGTATCCTATCTTATCATATAACTCG TTATGTTCTCATTTAGTGTCAATTATACCG CGGTACTGTAAATTC
hybridization sequence D′ hybridization sequence E′ hybridization sequence F′ STEM-hybrid′

(Stem 2) 5′-CGACGGCAGCGAGGC GAATTTACAGTACCG TATACGAAACAATCGTAAACATGAATCTAC TTTCCTTAACAATAGTACACATGTTAATGG AAAGTCATACTGTGATATACTAGTTTGTTG
bt-hybrid′ STEM-hybrid hybridization sequence A′ hybridization sequence B′ hybridization sequence C′

(Biotin-stem) 5′-GCCTCGCTGCCGTCG cca-biotin
bt-hybrid

(cy3-probe) 5′Cy3-cTAATGAAGA

(cy5-probe) 5′Cy5-tATGTAGATC

Supp. Fig. 2

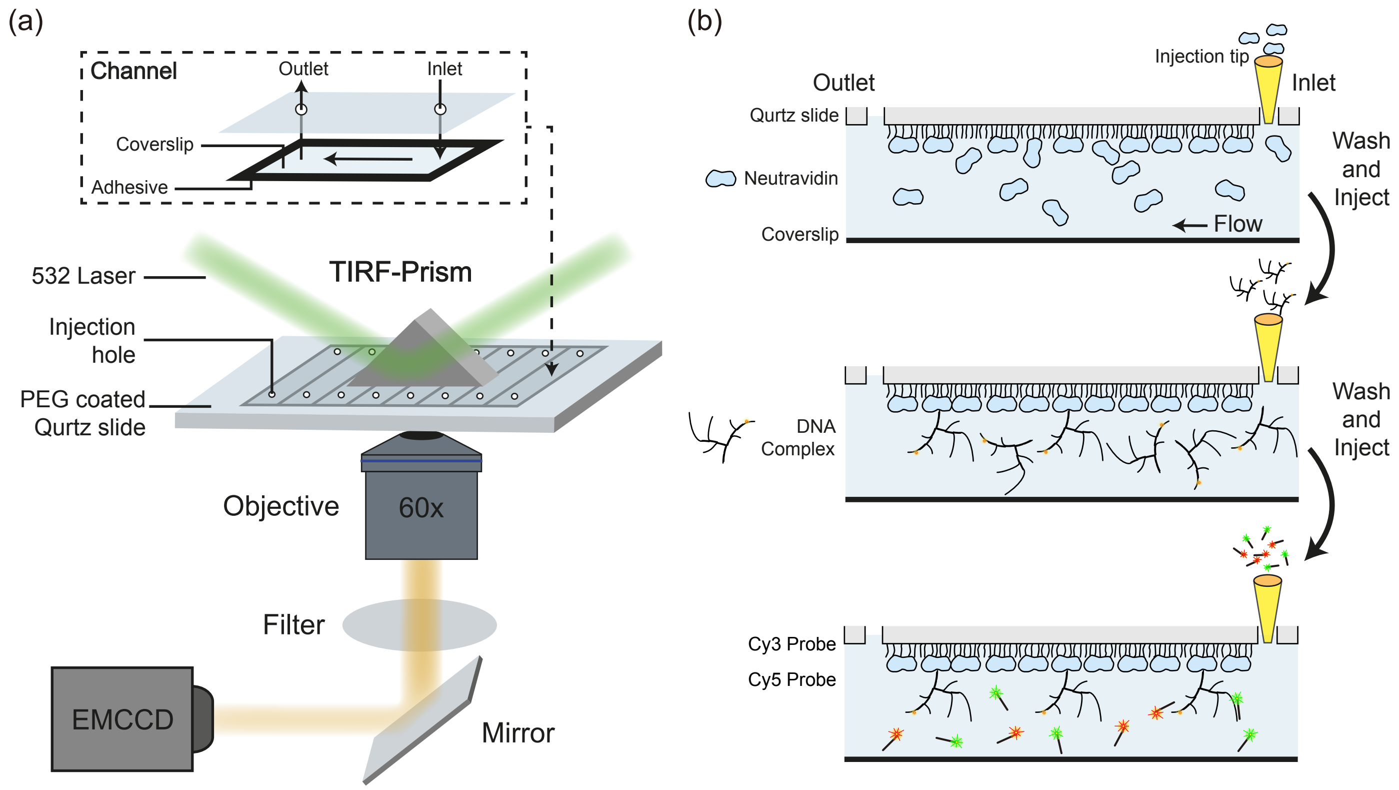

(a)
Channel
Outlet
Inlet
Coverslip
Adhesive
532 Laser
TIRF-Prism
Injection hole
PEG coated Qurtz slide
Objective
60x
Filter
EMCCD
Mirror
(b)
Injection tip
Outlet
Inlet
Qurtz slide
Neutravidin
Coverslip
Flow
Wash and Inject
DNA Complex
Wash and Inject
Cy3 Probe
Cy5 Probe

Supp. Fig. 3

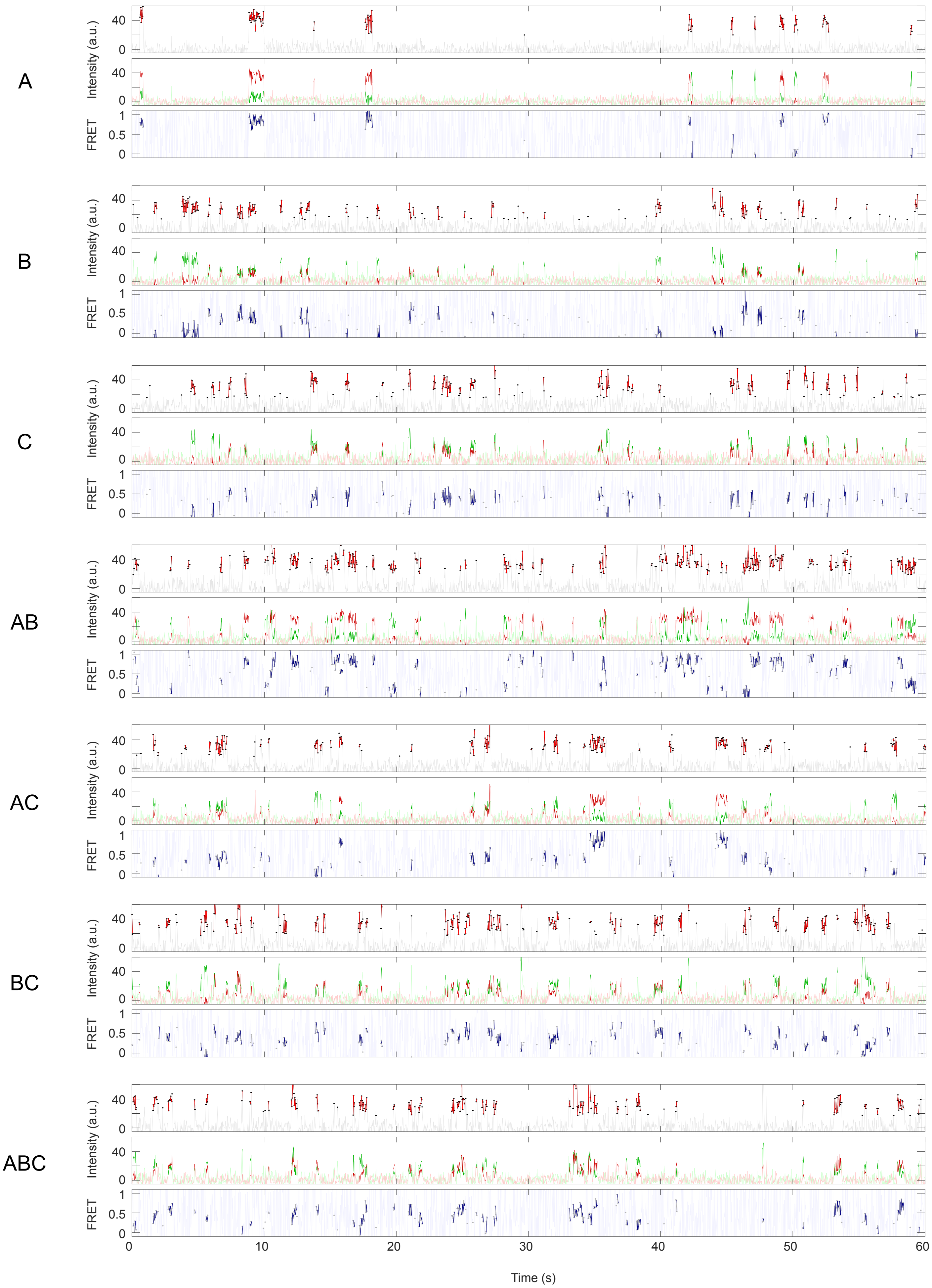

A
B
C
AB
AC
BC
ABC
Intensity (a.u.)
FRET
40
0
1
0.5
0
0
10
20
30
40
50
60
Time (s)

Supp. Fig. 4

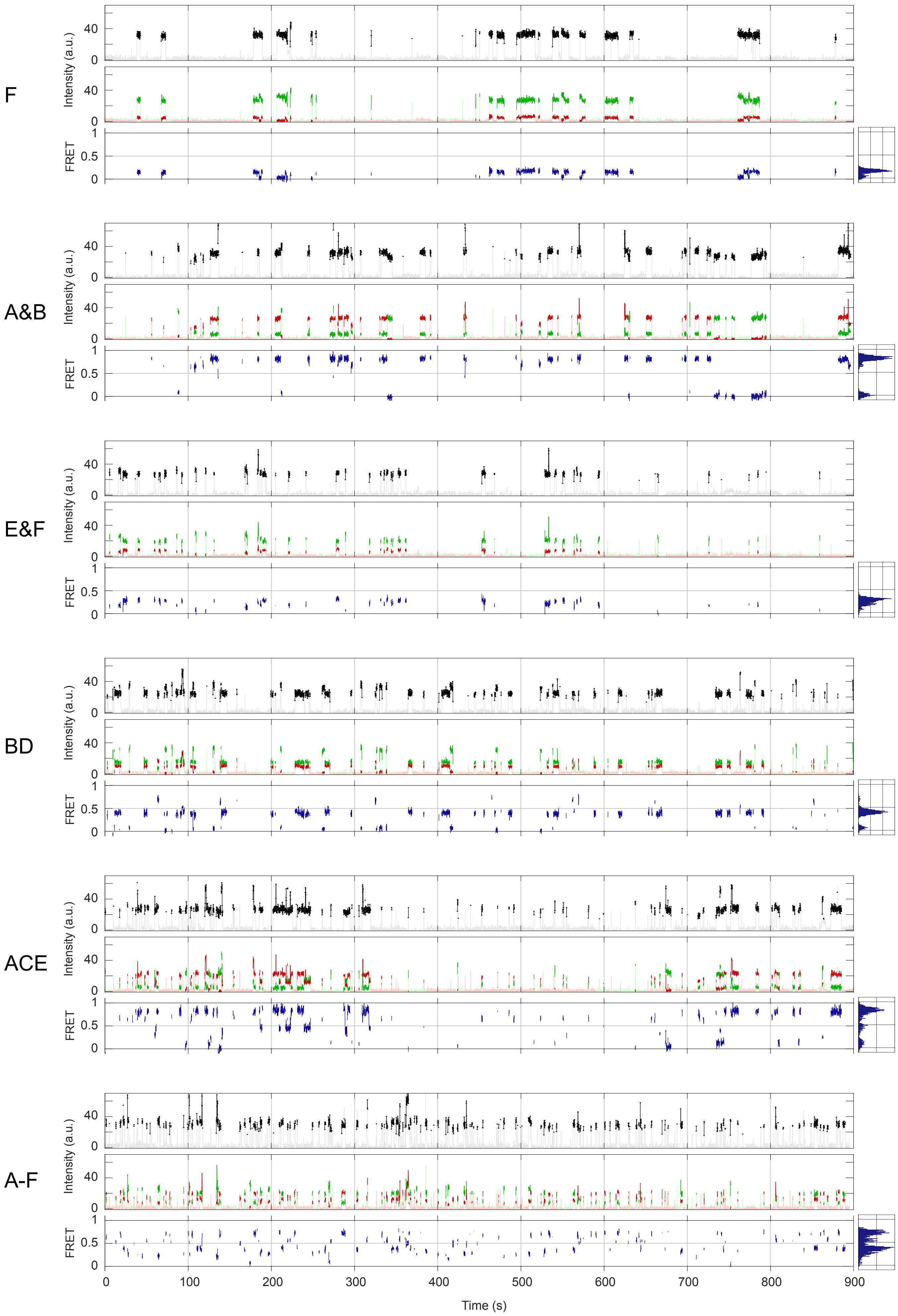

F
A&B
E&F
BD
ACE
A-F
Intensity (a.u.)
FRET
0
40
0.5
1
0
100
200
300
400
500
600
700
800
900
Time (s)
